# Towards an integrative multi-omics workflow

**DOI:** 10.1101/2021.07.26.453736

**Authors:** Florian Jeanneret, Stéphane Gazut

## Abstract

The advent of high-throughput techniques has greatly enhanced biological discovery. Last years, analysis of multi-omics data has taken the front seat to improve physiological understanding. Handling functional enrichment results from various biological data raises practical questions.

We propose an integrative workflow to better interpret biological process insights in a multi-omics approach applied to breast cancer data from The Cancer Genome Atlas (TCGA) related to Invasive Ductal Carcinoma (IDC) and Invasive Lobular Carcinoma (ILC). Pathway enrichment by Over Representation Analysis (ORA) and Gene Set Enrichment Analysis (GSEA) has been conducted with both features information from differential expression analysis or selected features from multi-block sPLS-DA methods. Then, comprehensive comparisons of enrichment results have been carried out by looking at classical enrichment analysis, probabilities pooling by Stouffer’s Z scores method and pathways clustering in biological themes.

Our work shows that ORA enrichment with selected sPLS-DA features and pathways probabilities pooling by Stouffer’s method lead to enrichment maps highly associated to physiological knowledge of IDC or ILC phenotypes, better than ORA and GSEA with differential expression driven features.

## 1 Introduction

Last years, high throughput biological technologies have greatly increased amount of data. Nowadays, public data repositories such as TCGA, METABRIC, CCLE and TARGET provide multiple types of omic data for the same samples[1]. A large number of multi-omics statistical data analysis has been developed recently taking several omic data blocks into account for patient profiling, clustering and biosignature detection[1].

Invasive Ductal carcinoma (IDC) and Invasive Lobular Carcinoma (ILC) patient data, from the TCGA consortium, are used in this work since they are phenotypes based on histological types exhibiting some molecular characteristics of breast cancer with many common disturbances expected[2].

Functional enrichment analysis is a very common method to reveal biological insights considering molecular differences between two phenotypes. It consists of presenting one list of significant pathways (e.g. from wikiPathways or Reactome) or ontologies (e.g. GeneOntology) potentially impaired between two sets of samples (e.g. sick and healthy patients). This method is based on a given list of genes that are differentially expressed and its information (e.g. p-value or log fold change). This Differential Expression Analysis (DEA) is implemented in some R packages such as DESEQ2, limma or edgeR. In our work, DESEQ2 and limma have been used. Nevertheless, differential expression and functional enrichment analysis are carried out for each block of omic data. Biologists and physiologists might assess sometimes contradictory enrichment results. Thereby, extracting biological signal in this situation where several blocks of omic data are available for each patient might be harder.

To the best of our knowledge, an integrative analysis to examine current potential of these methods towards functional enrichment of multi-omics data has not be addressed so far. Our work presents Stouffer’s pooling method and multi-block features selection benefits for altered biological pathways understanding in a multi-omics context.

## 2 Methods

RNAseq, RPPA (proteins) and clinical datasets from TCGA consortium were retrieved via firebrowse.org for RNAseq with 20 532 gene raw counts, RPPA data with 226 proteins and clinical data for 1097 samples. Only samples from primary tumor in common between three datasets have been kept for further analysis giving 573 samples for rna, protein and clinical data relative to ILC and IDC phenotypes.

Features with missing values and variance near to zero have been withdrawn. After this, datasets are constituted of 573 rows and respectively, for rna and protein data, of 19894 and 216 features.

To build functional enrichment analysis, duplicated features in RPPA data were averaged to give 173 features. Then, differential expression analysis by *DESeq2*[3] and *limma*[4] for respectively, RNAseq and protein RPPA data, have been computed and considered 8602 genes and 94 proteins as differential expressed features between IDC and ILC.

Biological knowledge database used in this work is Reactome[5], a free generalist database rich of thousands of human pathways.

Pathway enrichment analysis have been led by two types of statistical methods: Over Representation Analysis (ORA)[6] and Gene Set Enrichment Analysis (GSEA)[7].

ORA with differential expression features (DEF) is a statistical method to determine whether known biological processes (BP) are altered between experimental phenotypes. For a given biological process, it tests whether any difference of proportions that we observe is significant between number of DEFs in this BP relative to the number of the BP features and others DEFs regarding others features in database. This approach is equivalent to a one-sided proportion Fisher’s exact test.

GSEA is a widely functional enrichment approach that exploits a comprehensive list of features a metric ranking to sort list according to experimental phenotypes. Kolmogorov-Smirnov like statistic is computed and lets to conclude the putative alteration of each pathways in a database. Here, for RNAseq and protein RPPA data the ranking feature used were, respectively, statistic of *DESeq2* and log fold change from *limma*.

Stouffer’s probabilities pooling method is based on Z scores computed according to several p-values. Here, pathways p-values are obtained from enrichment analysis and each pathways shared by two omics enrichment results will have a computed Stouffer’s value. Fisher’s approach could be observed in such multi-omics context but Stouffer’s one is more reliable[8] and allows weights assignment for each data set. Weight in our case is the proportion of unique features recognised in pathways in enrichment results, regardless significance, out of total number of features in this omic data block.

For the *i^th^* p-value, where Φ is standard normal cumulative distribution, we have:

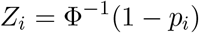

Then, for *k* omic datasets we compute pooled Z score:

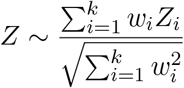

Finally, probability after fusion *p_z_* is:

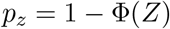

To determine small subsets of biological features to enrich, sparse Partial Least Squares-Discriminant Analysis (sPLS-DA)[9] from the R package *mixOmics* has been used. sPLS-DA approach identifies highly correlated multi-omics signature between blocks and linked to phenotypes. Here, this statistical method computes a classification model based on RNAseq and RPPA data toward IDC and ILC phenotypes classification. This model is fitted by 80% of total number of samples (458 out of 573 samples) and forecasting performances is evaluated on external dataset of 20% (115 samples) remaining data. The number of selected features per component has been assessed by cross-validation (10 folds, repeated 20 times).

Enrichment results from ORA or GSEA analysis may be hard to interpret with the classical table output format when the number of significant pathways is large. The EnrichmentMap Cytoscape module[10] builds pathways network where nodes are pathways and bond thicknesses depend on proportion of genes shared between two pathways. This visualisation, helped by AutoAnnotate module as seen in [11] that computes network clusters, lets to detect biological themes for which more than one pathways are found in enrichment results and to be focused on larger ones.

All selected features from upstream analysis have been converted to entrez id for enrichment analysis.

## 3 Results

To assess our multi-omics workflow analysis (fig. **1**), ORA and GSEA enrichments of Reactome pathway database were conducted.

**Fig. 1.**
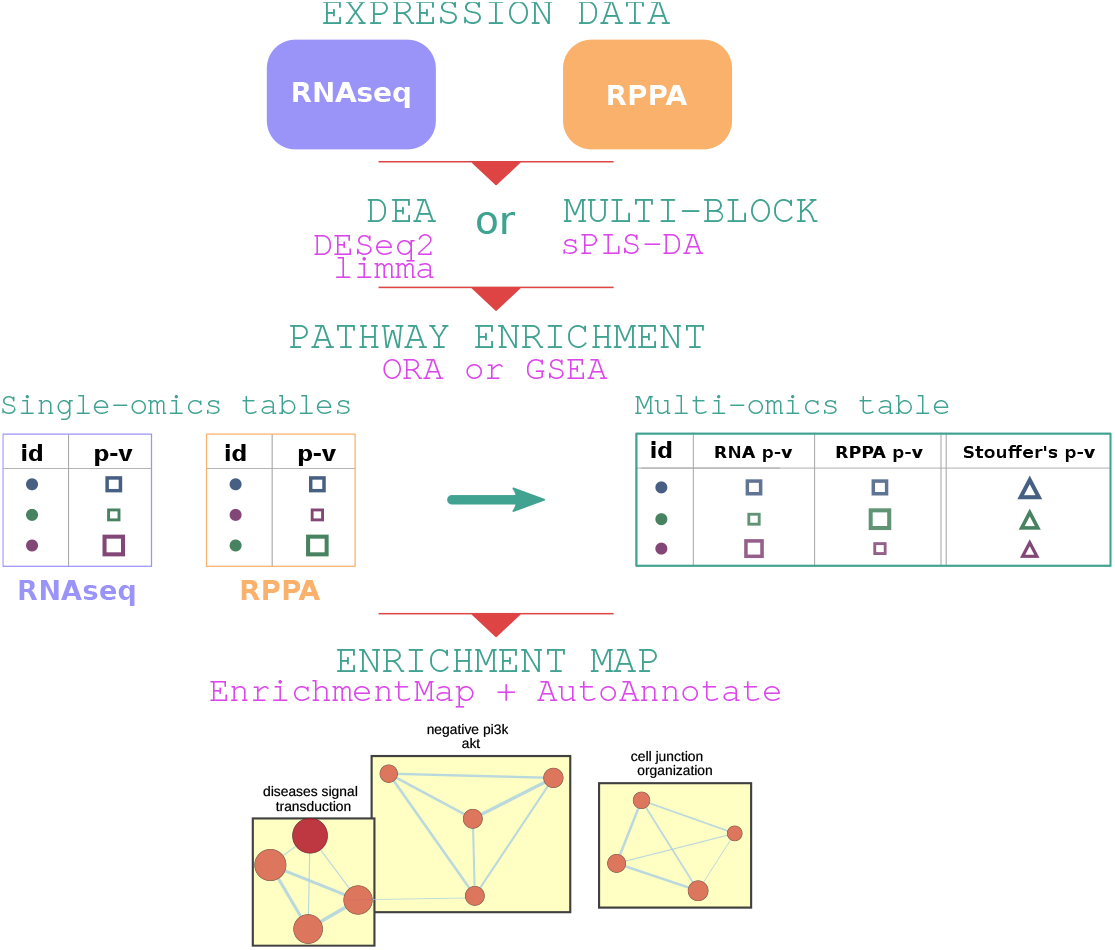
Multi-omics analysis workflow. Expression data for several biological actor (e.g. RNAseq and RPPA) and for the same patients begets a multi-omics context. To reveal altered pathways between patient phenotypes we are aiming at enrichment analysis by ORA or GSEA. Before this step, Differential Expression Analysis (DEA) is carried out for classical ORA and GSEA by DESeq2 and limma analysis or features are selected by sPLS-DA for a multi-omics driven ORA. Then, each enrichment results from omic blocks are merged by p-values using Stouffer’s probabilities pooling method. All these enrichment results are visualised by enrichment map to interpret easily highlighted biological themes.

Firstly, towards ORA and GSEA enrichment, Differential Expression Analysis (DEA) has been computed on two omic datasets RNA and RPPA. For ORA and for each omic data, a list of significant features differently expressed between IDC and ILC is saved for feature p-values under 0.05 after Benjamini-Hochberg correction. Then, features have been selected by sPLS-DA multi-block model to compare biological insights with previous classical methods.

Secondly, classical functional enrichment analysis ORA and GSEA have been carried out for each set of omic features. ORA was fed by selected features and GSEA by all features ranked by statistic or log fold computed by DEA (see *Methods*).

In addition, for a given pathway shared in RNAseq and RPPA enrichment results, Stouffer’s probabilities pooling method has been used to merge p-values and build a multi-omics table with significant pathways harnessing multi-omics information. Finally, for each enrichment result (e.g. ORA with RNAseq selected features, multi-omics table with pathways and Stouffer’s p-values) an enrichment map has been drawn to visualise significant biological themes and to compare classical and multi-omics analysis with Stouffer’s probabilities pooling and sPLS-DA selected features methods.

### 3.1 Over Representation analysis (ORA) of DESeq2/limma features

The first functional analysis used is ORA on the two sets of features obtained by Differential Expression analysis. It consists in detecting the enriched, and so potentially altered, pathways by Fisher’s exact test (see *Methods*).

In the first hand for RNAseq data, with 8602 selected genes after *DESeq2* Differential Expression analysis, 336 pathways have been detected as significantly enriched under 0.05 p-value threshold after Benjamini-Hochberg multiple testing corrections (tab.**1**). To get a better sight of enrichment results, *EnrichmentMap* Cytoscape module has been used in this case and further enrichment results (e.g. GSEA) and for each omic dataset. Once enrichment map built, pathways clusters have been computed and could be seen as biological themes.

**Tab. 1.** Number of significant pathways (out of all pathways with at least one gene enriched) according to features selection method and enrichment results. “*DESeq2/limma*” means features were selected based on differential expression analysis and probabilities under 0.05 threshold. Multi-omics table presents a Stouffer’s pooled p-value for each pathway shared by RNAseq and RPPA enrichment results. Statistical significance is defined, once Benjamini-Hochberg multiple testing correction has been applied, by each pathway probability below the 0.05 threshold.

|  | RNA | Protein | multi-omics table |
| --- | --- | --- | --- |
| ORA (*DESeq2/limma*) | 336 (1827) | 226 (788) | 118 (1827) |
| GSEA (*DESeq2/limma*) | 150 (1473) | 0 (109) | 14 (1477) |
| ORA (*sPLS-DA*) | 5 (280) | 128 (522) | 47 (653) |

On this enrichment map drawn with significant pathways of this ORA, we observe 41 functional themes. The largest clusters with respectively, 95 and 45 pathways, are known in breast cancer: *regulation of mitotic cycle* and *NOTCH signaling*[12]. In fact, a lot of significant pathways in ORA are very general human pathways deregulated in cancers, leaving biological insights relative to ILC and IDC phenotype hard to discover and interpret in this enrichment map.

On the other hand, protein data gives similar results with 226 pathways under statistical threshold after ORA of 93 features (tab.**1**). For instance, 42 clusters in enrichment map with 35 pathways in *overexpressed ERBB2*, 16 pathways in *regulation of mitotic cycle*, 11 pathways in *moderate kinase BRAF*, 10 pathways in *MAPK, PI3K/Akt signaling* and 8 pathways for *RB1 signaling* clusters are related to breast cancer or cancer biology in literature[13,14]. They are not very specific to ILC and IDC phenotypes according to our knowledge except *overexpressed ERBB2* with key ERBB2 gene known to be expressed differently between ILC and IDC samples[15]. Others biological clusters are made up of 7 to 2 pathways, for a mean of 3.5 nodes by clusters. Thus, these clusters are not discussed with assumption that little clusters are more susceptible to be false positive ones.

In this case, 43 pathways are in common between these two enrichment result tables from RNAseq and RPPA (protein) features. We may think overlapped significant pathways between several omic data is a good point to start biological interpretation but it would miss not significant pathways in several data block could be a relevant information. For instance, 2 pathways could have both 0.06 of significance. Classical analysis would reject these pathways for each block.

To get a better biological vision taking into account statistical information shared in several omic blocks and to explain altered pathways between complex phenotypes, we propose here Stouffer’s pooling method (see *Methods*) as seen in different multi-omics tools[16,17]. Therefore, pathways in common between RNAseq and protein enrichment results lead to a multi-omics enrichment table where best pathways are those with best probabilities taking two omic datasets into account.

We observe 118 pathways according to Stouffer’s value under the statistical 0.05 threshold (tab.**1**). Thus, 85 more pathways are significant compared to retrieving only overlapped pathways between significant enrichment results of RNAseq and protein data. Enrichment map obtained by these Stouffer’s values consists in 20 clusters and helps to remark *regulation of mitotic cycle* theme with *g1 transition degradation*, *cycle anaphase mitotic*, *MAPK signaling* theme with *diseases signal transduction* and *mitotic G2 phase* clusters with respectively, 46, 8, 5 and 4 pathways. These biological themes are also relative to breast cancer[14] but not very specific to ILC or IDC biology. Other clusters are 3 or 2 nodes driven, among them *Rho GTPases signaling* could be interesting according to[18] but *hiv life cycle* is also present and seems to be false positive biological theme. We prefer not to comment these little clusters regarding this statement and high number of pathways found significant.

Great amount of pathways as seen in this section may be only false positive ones due to statistical hypothesis and method used. To address this, GSEA was carried out on same datasets and with same analysis procedure to compare new enrichment results with Stouffer’s values and enrichment maps.

### 3.2 Gene Set Enrichment Analysis (GSEA) of DESeq2/limma information

GSEA has been undertaken as ORA with Reactome database with significant RNAseq and protein features given by Differential Expression analysis (see *Methods*). Firstly, 145 pathways are considered significant according to adjusted p-values from RNAseq features functional analysis (tab.**1**). Among them, after enrichment map clustering, we find two major themes: 96 pathways are clustered in *regulation of mitotic cell cycle* and 16 in *Rho GTPases signaling*. These two biological sets are correlated to breast cancer where revamped *Rho GTPases* pathway plays a critical role in mitotic cell cycle, cellular transformation and cancer cell proliferation[18]. More, RhoA seems to be associated with collective cell invasion in IDC subtype carcinoma[19].

Secondly, protein features GSEA has no signal: 0 significant pathways have been found significant (tab.**1**). The nature of the data, two subtypes of the same disease, could explained protein expression differences too tiny to be distinguished. This case, where overlap between significant pathways from RNAseq and RPPA features GSEAs is of course null, illustrates weakness to look only significant pathways in common between several enrichment results obtained by several omic data.

Thereby, to handle multi-omics interpretation, a multi-omics enrichment table based on Stouffer’s probabilities merging both adjusted p-values from GSEAs was calculated. This multi-omics table presents 15 significant pathways under 0.05 Stouffer’s value threshold (tab.**1**) leading biological interpretation possible where both omic data are taken in account whereas classical enrichment results from GSEA of protein features gives no signal.

Enrichment maps obtained by RNA features enrichment table and multi-omics table are quite different with 150 nodes and 16 respectively. Enrichment map based on multi-omics table enables to visualise pathways sharing features in common. We can observe two previous themes about *mitotic cell cycle* and *rho GTPases signaling* with 13 and 2 pathways respectively, giving a easier biological knowledge graph to interpret. As seen with enrichment map from rna features GSEA, these two biological themes are related to complex biological phenotypes ILC and IDC carcinomas[18,19]. The use of Stouffer’s method has reinforced interpretation given by RNAseq analysis with a multi-omics insight.

Classical enrichment results, using DEA to feed ORA and GSEA, have shown pathways related to breast cancer and one or two biological themes really associated to ILC or IDC carcinomas. Multi-omics tables based on Stouffer’s probabilities pooling method has led to more targeted and confident biological interpretation taking both enrichment results into account from each kind of omic features. It is now a question of completing the multi-omics approach by assessing a multi-omics variable selection method substituting the DEA.

### 3.3 ORA of sPLS-DA features

Last but not least, multi-omics as a research theme is rich in statistical methods developed or adapted toward better biological understanding[1]. Some of methods harnessing discriminant features and correlations at the same time (e.g. sPLS-DA[9], RGGCA[20]). As such, sPLS-DA from mixOmics R package has been used for further analysis. sPLS-DA is dedicated to select features from several numerical matrices with discriminant power and correlated between datasets and phenotypes. Here we wish assess enrichment tables and enrichment map computed with these multi-omics selected features and how this kind of method could lead to a better biological understanding.

Selected features of each latent variables given by sPLS-DA model and for each omic dataset are stored. 49 genes and 39 proteins have been selected in this process. More, to evaluate specificity of these features toward ILC and IDC carcinomas phenotypes, phenotype forecasting performances has been evaluated. An accuracy of 76% has been obtained with test on external dataset of RNAseq and RPPA data. Thus, these selected features are considered specific to the biological problem. Therefore, ORA enrichment analysis has been carried out for rna and protein features to detect putative altered biological pathways. ORA enrichment results has given 5 and 128 pathways for, respectively, rna and protein features selected by sPLS-DA. Relative enrichment maps computed by EnrichmentMap Cytoscape module present these pathways as biological themes in network clusters.

Firstly, with 49 rna features, we detect 5 significant pathways (tab.**1**) and only one biological cluster by enrichment map with 3 pathways. This one describes *signaling nuclear receptors* theme with *ESR-mediated signaling* pathways known to be altered in breast cancer namely with ESR gene mutation[14,15].

Secondly, 128 significant pathways have been found by ORA with 39 selected proteins (tab.**1**). Same as before, enrichment map has been drawn to visualise biological themes. Among 29 biological clusters, some of them are especially correlated to ILC and IDC carcinomas literature. Indeed, *Tie2 overexpressed ERBB2* (16 pathways) and *AKT network PI3K* (5 pathways) biological themes seen in protein features ORA, *regulation signaling kit* (6 pathways) and *extra nuclear ESR* (3 pathways) have central mutated genes and differently expressed between ILC and IDC subtypes[14,15].

More, *cell-cell junction organisation* pathways are highlighted in a cluster. The latter is directly consistent with the known biology of ILC carcinoma characterised by lack of CDH1 expression driving dysregulation of cell-cell adhesion[21] seen previously in rna features ORA as an ILC characteristic. Others clusters as *kinase BRAF mutants* (10 pathways) or *MAPK signaling* (4 pathways) detected in ORA and *activated downstream FGFR3* with FGRF1, FGFR2 and FGFR3 signaling pathways could be linked to general breast cancer biology[13,14].

To reinforce insights discovery from several type of omic data enrichment results, Stouffer’s probabilities pooling method have been used as before for each pathways corrected p-value shared between enrichment results.

Only 3 pathways with corrected p-values under 0.05 threshold are shared between two sets of rna and protein enrichment results by sPLS-DA features selection. Once again, this shows that considering only pathways overlap from enrichment results in a multi-omics context is too conservative.

Thereby, to merge p-values from enrichment tables for same pathways, Stouffer’s values are calculated and 47 significant pathways are found (tab.**1**). Enrichment map drawn with multi-omics table (fig.**2**) has built 11 biological clusters whose several directly related to phenotypes: *Tie2 overexpressed ERBB2*, *negative PI3K AKT*, *cell junction organization* and *estrogen dependent expression* clusters built with, respectively, 10, 4, 4 and 4 pathways. In fact, we know ERBB2 gene expression is different between ILC and IDC samples[15], PI3K is a mutated gene with single nucleotide variation and at high expression level in particular in IDC subtype[2,14], 55–72% of IDCs being estrogen receptor positive compared with 70–92% of ILCs[15] and *cell-cell junction organization* theme is directly consistent with the known biology of ILC carcinoma with dysregulation of cell-cell adhesion[21]. The latter is retrieved only with sPLS-DA features enrichment, not with ORA and GSEA computed on DESeq2 and limma features information. We found also breast cancer related biological clusters such as *MAPK signaling* cluster with 4 pathways.

**Fig. 2.**
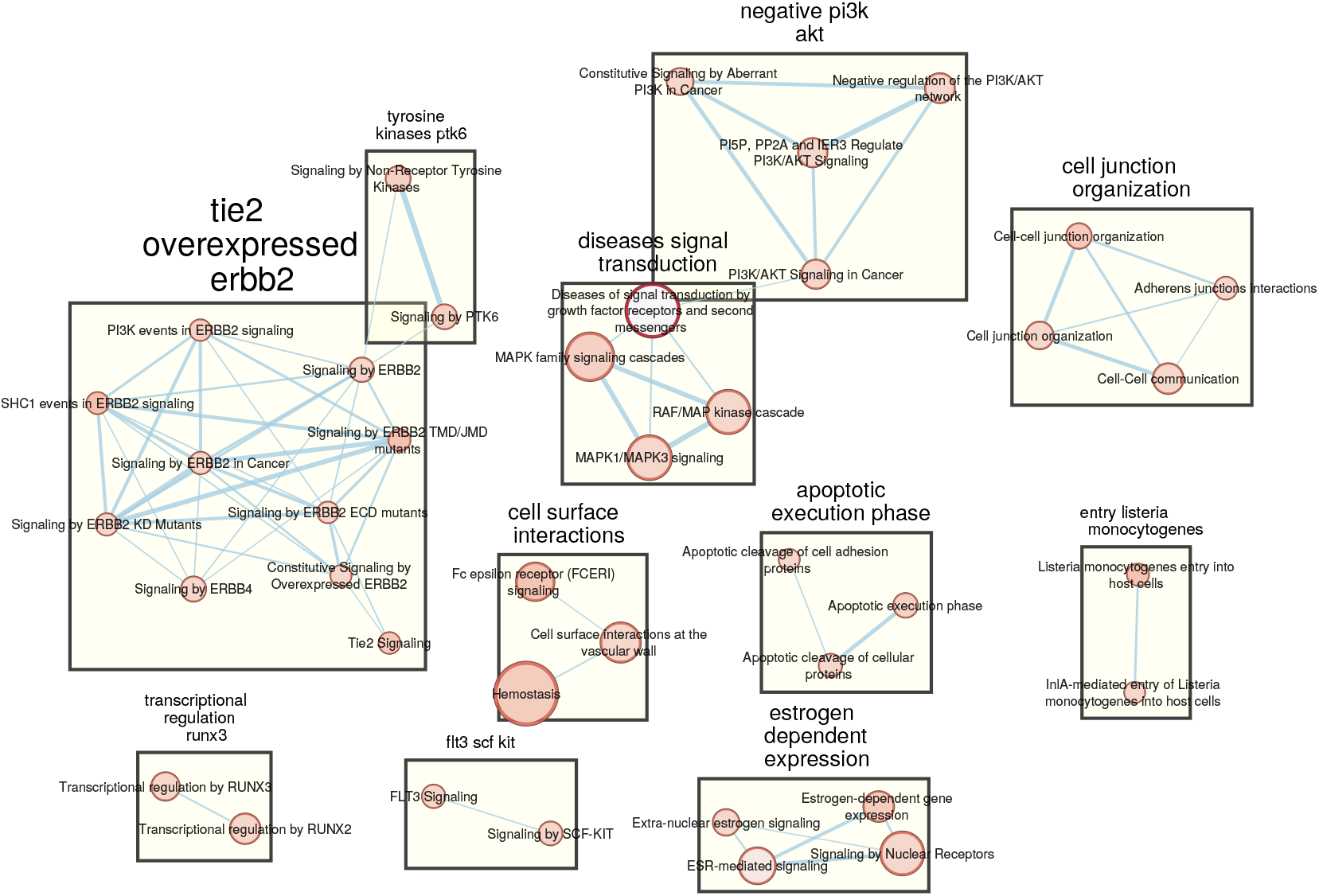
Enrichment Map of significant pathways obtained by features selected by sPLS-DA method. Statistical significance is defined according to Stouffer’s value under the 0.05 threshold calculated for each pathways shared by RNAseq and RPPA enrichment results and corrected according to Benjamini-Hochberg method.

Thereby, sPLS-DA driven feature selection enables a more phenotype-targeted functional enrichment. Based on Reactome database, biological themes associated to ILC and IDC carcinomas differences are highlighted in enrichment maps after pathways clustering. Moreover, Stouffer’s probabilities pooling method shows great benefits to more easily interpret several enrichment results in a multi-omics context.

### 4 Discussion

To boil down results, using Stouffer’s probabilities pooling method, to build multi-omics enrichment table, leads to make easier biological interpretation between several enrichment analysis carried out on omic datasets. Moreover, using multi-omics features selection by sPLS-DA reveals better biological insights associated to complex histological phenotypes than classical functional analysis such as ORA and GSEA fed by DEA information.

Nevertheless, some points could be explored to reinforce these statements. Firstly, carrying out same analysis on other datasets from TCGA with sufficient number of common samples between several kind of omics and by adding other biological actors as microRNAs or methylation data.

Secondly, we may check results with other databases. For instance, other related to pathways (e.g. KEGG, wikiPathways), Gene Ontology to investigate molecular functions, or more detailed about biological domains (PANTHER, MSigDB). Moreover, some solutions to merge generalist pathways databases have shown good results, and it would be interesting to implement in this workflow, such as MPath[22] where KEGG, Reactome and wikiPathways are merged toward better covering of biological pathways.

Finally, sPLS-DA is one a plenty of multi-omics statistical analysis developed last years[1]. In fact, it could be also interesting to investigate this workflow with other unsupervised or supervised multi-block methods.

## 5 Conclusion

We present assessing of our multi-omics workflow based on sPLS-DA selected features, enriched pathways probabilities pooling and enrichment mapping. We show good results towards the efficient management of functional enrichment results in a multi-omics approach. Method considering significant pathways overlap between several omics enrichment results is too conservative and does not take account advantage of multi-omics context whereas Stouffer’s probabilities method leads to more efficient pathways highlighting. Moreover, features information from differential expression analysis tends to enrich pathways associated to breast cancers but not very specific to IDC or ILC. In contrasts, pathways enriched by sPLS-DA features are clustered in biological themes highly related to these histological types.

To conclude, pathways enrichment results of sPLS-DA features pooled by Stouffer’s method seems to lead to better biological processes discovery in a multi-omics context.

